# Incorporating Prior Knowledge into Regularized Regression

**DOI:** 10.1101/2020.03.04.971408

**Authors:** Chubing Zeng, Duncan Campbell Thomas, Juan Pablo Lewinger

## Abstract

**Motivation:** Associated with genomic features like gene expression, methylation, and genotypes, used in statistical modeling of health outcomes, there is a rich set of meta-features like functional annotations, pathway information, and knowledge from previous studies, that can be used post-hoc to facilitate the interpretation of a model. However, using this meta-feature information a-priori rather than post-hoc can yield improved prediction performance as well as enhanced model interpretation.

**Results:** We propose a new penalized regression approach that allows a-priori integration of external meta-features. The method extends LASSO regression by incorporating individualized penalty parameters for each regression coefficient. The penalty parameters are in turn modeled as a log-linear function of the meta-features and are estimated from the data using an approximate empirical Bayes approach. Optimization of the marginal likelihood on which the empirical Bayes estimation is based is performed using a fast and stable majorization-minimization procedure. Through simulations, we show that the proposed regression with individualized penalties can outperform the standard LASSO in terms of both parameters estimation and prediction performance when the external data is informative. We further demonstrate our approach with applications to gene expression studies of bone density and breast cancer.

**Availability and implementation:** The methods have been implemented in the R package *xtune* freely available for download from CRAN.

## 1 Introduction

Predicting outcomes based on genomic biomarkers such as gene expression, methylation, and genotypes is becoming increasingly important for individualized risk assessment and treatment (Kamel and Al-Amodi, 2017). As an example consider predicting mortality from breast cancer after surgical treatment based on gene expression profiles (Nuyten *et al.*, 2006). Since genomic studies typically have more available features than subjects, a common approach to develop prediction models based on genomic features is to use regularized regression methods, which can handle high-dimensional data. In addition to regularization, sparsity inducing regression approaches such as the LASSO can also perform feature selection. In the context of genomic studies, feature selection is critical for yielding interpretable models that provide insight into potential biological mechanisms and which can, in turn, facilitate adoption by practitioners. To enhance model interpretability, it is common to examine features selected in a model in relation to available information about gene function and previous studies. For example, analyses can be conducted to formally assess whether the selected features are enriched in particular metabolic pathways or gene ontology annotations. This kind of post-hoc analysis relating genomic features to existing knowledge about them, hereafter referred to as genomic meta-features, can provide valuable biological insight and validation for a prediction model. In this paper, we propose a new approach that exploits genomic meta-features a-priori rather than post-hoc, to improve prediction performance and feature selection, and to enhance the interpretability of models developed using penalized regression.

Most penalized regression methods apply a single global penalty parameter to all regression coefficients, effectively treating all features or predictors equally in the model building process. This can result in over-shrinking of important coefficients and under-shrinking of unimportant ones, with a corresponding loss in prediction ability. In genomics and other applications, there is often a great deal of prior knowledge about the features. Examples of prior knowledge include (1) gene function annotation from databases like the Gene Ontology Project (Ashburner *et al.*, 2000); (2) gene-disease co-occurrence scores from text-mining biomedical abstracts (Pletscher-Frankild *et al.*, 2014; Rouillard *et al.*, 2016); (3) deleterious somatic mutations in the Catalogue of Somatic Mutations in Cancer (COSMIC)(Forbes *et al.*, 2010). With the main goal of improving prediction but also to improve feature selection performance, parameter estimation, and model interpretability, we extend the standard LASSO regression to allow for penalty terms that depend on such external meta-features. Specifically, rather than using a single penalty parameter that controls the global amount of shrinkage across all regression coefficients, our model allows each coefficient to have its own individual penalty parameter, which is in turn modeled as a function of the meta-features. We focus on the LASSO penalty because of its widespread use but address potential extensions to other penalties in the discussion.

Previously, other variants of LASSO regression have been introduced to allow either coefficient-specific penalties or multiple tuning parameters. Yuan and Lin (2006) proposes group LASSO that extends LASSO to grouped predictors. Zou (2006) adjusts the penalty differently for each coefficient by using a vector of adaptive weights. The group Lasso applies only to grouping variables and the adaptive Lasso uses pre-specified weights obtained from the initial estimate of the coefficients using the same data as the data used for regression. Neither of these approaches incorporate a general set of meta-features. Boulesteix *et al.* (2017) proposed the integrative LASSO with penalty factors method that assigns different penalty factors to different data modalities such as gene expression, methylation, and copy number. They use cross-validation to choose the penalty parameters based on prediction performance. In practice, the number of different modalities they can allow is up to 4, due to computational bottle-neck.

In addition to those above, several other methods have been proposed previously to make use of prior knowledge of the features. Tharmaratnam *et al.* (2016) suggests using biologic knowledge to derive a set of features that could replace the genes set by chosen by LASSO with minimal loss in predictive power. However, ‘experts’ are required to assess the importance of each gene. Tai and Pan (2007) partitions the features into groups and shrink the features of different groups by different magnitudes. Shrinkage for groups is considered fixed and arbitrary but is not data-dependent and the use of an external dataset only provides information on the grouping of predictors. van de Wiel *et al.* (2016) proposes an adaptive group-regularized ridge regression that account for group structure as the group LASSO and allow group-specific penalties. However, the method require partitioning of the features into different groups, and the external information is only used for guiding the partition. The approach proposed by Bergersen *et al.* (2011) is also in the form of the adaptive Lasso. Instead of constructing weights from the same data *X* and *Y*, they use the Spearman correlation coefficients or the ridge regression coefficients between the features *X* and external information as the penalty weights.

Our approach is distinguished from these methods in that (i) we adopt an empirical Bayes approach to estimate the hyper-parameters instead of cross-validation, which allows us to estimate feature-specific tuning parameters; (ii) the magnitude of the penalty terms are modeled as a log-linear function of the external information and are estimated from a ‘second-level’ regression; (iii) our approach is not restricted to meta-features that define feature groupings but can handle meta-features of any type including quantitative ones.

## 2 Methods

### 2.1 Random effect formulation of the LASSO

We start by considering a standard linear regression model *Y* = *X**β*** + ***ϵ***, where *Y* is the vector of observed responses for *n* subjects, *X* is an *n* × *p* matrix of genomic features, and ***ϵ*** represents independent errors with zero expectation and a common variance *σ*^2^. The LASSO regression shrinks the regression coefficients by imposing a *L*_1_ penalty on the sum of absolute value of regression coefficients (Tibshirani, 1996). The objective function of LASSO is:

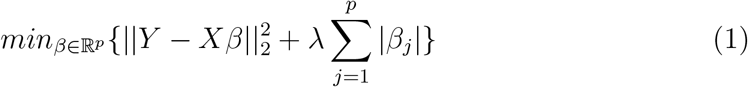

Most penalized regression approaches have a Bayesian interpretation. The LASSO regression can be equivalently formulated as a Bayesian or random effects model where the coefficients are modeled with a double exponential (a.k.a. Laplace) prior distribution condition on *σ*^2^ (Tibshirani, 1996). In specific, we assume that the distribution of the response variable *Y* conditional on the regression coefficients ***β*** and *σ*^2^ follows a normal distribution. The LASSO coefficient estimates that solves (1) can be equivalently characterized as the posterior mode, or maximum a posteriori (MAP) in a Bayesian model with the following double exponential prior distribution for ***β*** conditional on *σ*^2^:

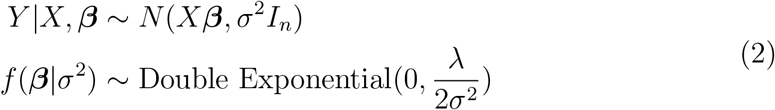

Our model extends the objective function of LASSO to allow each coefficient to have its own individual penalty parameter, which is in turn modeled as a log-linear function of the meta-features. The objective function of external information tuned (xtune) LASSO is:

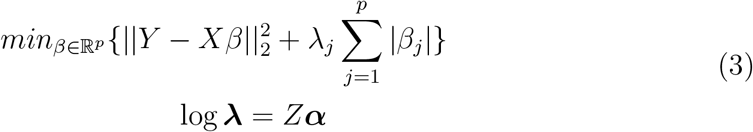

where *Z* is the meta-feature of dimension *p* × *q*, the *j^th^* column of *Z* comprise information about feature *j*. ***α*** is the unknown hyperparameter vector that links the individual penalties and the external information.

By using the Bayesian formulation on penalized regression, external covariates can be incorporated via the prior on the model coefficients. Here we allow differential penalization by assigning priors with different scale parameters to regression coefficients. In specific, the double exponential random effect formulation of the standard LASSO in (2) assumes a single common ‘prior’ variance for all *β*_*j*_’s. In our extension, we let each *β*_*j*_ have a double exponential random effect distribution but with its own individual hyperparameter *λ*_*j*_, where the hyperparameter vector ***λ*** is in turn modeled as a log-linear function of the external information available for feature *j*, *Z*.

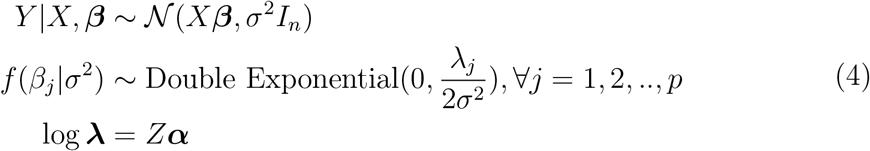

The maximum a posteriori (MAP) estimator under the random effects model in (4), which is conditional on *σ*^2^, is equivalent to the estimator minimizing the LASSO objective function with external information driven penalization in (3). The population variance parameter *σ*^2^ can estimated from the data or given a point-mass prior (Li and Lin, 2010). In this paper, we estimate *σ*^2^ using the method proposed by Reid *et al.* (2013) and assume it is ‘set in advance’. Park and Casella (2008) and Li and Lin (2010) used a specification that assigns a non-informative prior for *σ*^2^ and used a Gibbs sampler based on full conditional distributions to obtain regression coefficients estimates. Our method is different in that, rather than extending the model to include Bayesian inference over the hyperparameters, we use a empirical Bayes approach that maximize the log likelihood of hyperparameters. That is, the hyperparameters ***α*** are estimated from the data by first marginalizing over the coefficients *β* and then performing what is commonly referred to as empirical Bayes, evidence maximization or type-II maximum likelihood Tipping (2001).

### 2.2 Empirical Bayes parameter tuning

Unlike the standard LASSO, the proposed model has a potentially large number of hyperparameters (***α***_1×*q*_), so tuning them by cross-validation is not feasible. Instead, we propose an empirical Bayes approach for directly estimating the hyperparameters based on the training data. Specifically, the empirical Bayes of ***α*** (hence ***λ***) is obtained by maximizing the marginal likelihood calculated by integrating out the random effects ***β*** from the joint distribution of *Y* and ***β***.

The marginal likelihood of ***α*** is given by:

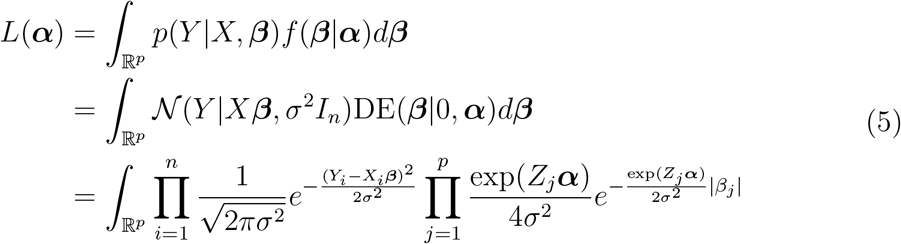

When X is not orthogonal, the marginal likelihood resulting from the *p*-dimensional integral in (5) does not have a usable close-form solution. Foster *et al.* (2008) proposes using a Laplace approximation to the integral which has simple close-form solution. Motivated by their approach, we propose a simpler new approximation which uses a normal distribution with the same prior variance to approximate the double exponential distribution. That is, we use a normal distribution 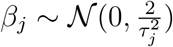 to approximate the double exponential distribution *β*_*j*_ ~ Double Exponential(*τ*_*i*_), yields closed form solution for the approximated marginal likelihood:

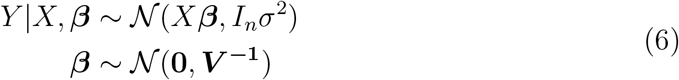

where

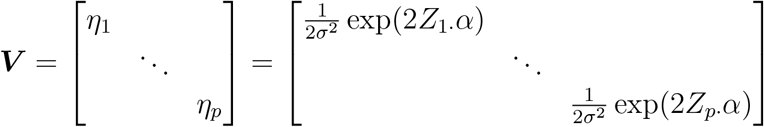

Therefore, the approximate log marginal likelihood of ***α*** integrating out ***β*** is:

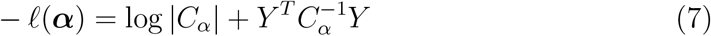

where *C*_*α*_ = *σ*^2^*I*+*XV*^−1^*X*^*T*^. The approximated log likelihood (7) is then maximized to obtain ***α*** estimates. Once ***α*** known, hence the penalty parameters vector ***λ*** known, the *glmnet* package (Jerome Friedman, Trevor Hastie and Tibshirani, 2010) in the R language is used to implement the LASSO with given penalty vector.

### 2.3 Optimizing the Objective Function

The objective function (7) is non-convex, making it intrinsically a challenging problem. We note that the approximated model (6) is closely related to the model specification of the Automatic Relevance Determination (ARD) (MacKay, 1992; Neal, 1995; Tipping, 2001) method widely used in the field of signal processing. Wipf and Nagarajan (2008, 2014) describe a Majorization Minimization (MM) procedure that uses a reformulation of ARD to optimize the non-convex optimization function by solving a series of easier re-weighted *L*_1_ problem. Motivated by their idea, we propose an iterative re-weighted *L*_2_ optimization algorithm described in detail below. Note that this non-convex optimization problem is a special case of the difference of convex functions (DC) problem (Le Thi and Pham Dinh, 2018).

Note that the log-determinate term log |*C*_*α*_| is a concave function in ***α*** (Boyd and Vandenberghe, 2004). A majorization function of log |*C*_*α*_| is its slope at the current value *α* of log |*C*_*α*_|:

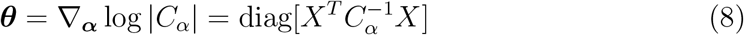

The 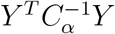 term in (7) is convex. Therefore, at current value of ***α***, the majorization function for −*ℓ*(***α***) is 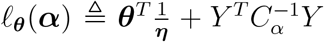. Given ***θ***, ***α*** is updated by:

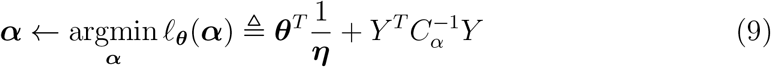

Although the objective function (9) is convex, it is slow to optimize in practice. We use one more MM procedure for optimizing (9). Note that the data dependent term 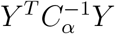 can be re-expressed as:

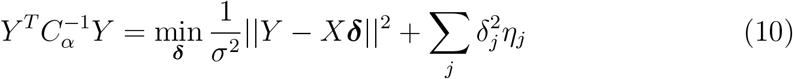

We therefore introduce another auxiliary term *δ*, the upper-bounding auxiliary function for *ℓ*_***θ***_(***α***) is:

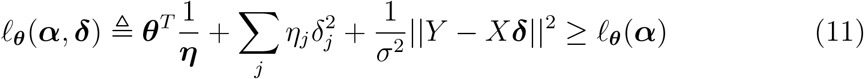

The ***α*** value that minimizes equation (9) can be estimated by iteratively updating ***δ*** and ***α*** in equation (11). For any ***δ***, ***α*** is estimated by minimizing

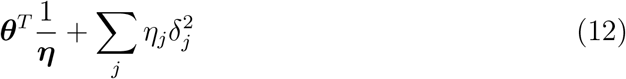

Given ***α***, ***δ*** is updated by:

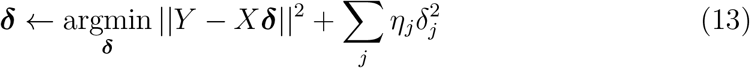

Equation (13) has a weighted convex *L*_2_ regularized cost function, and it can be optimized efficiently using *glmnet*. In summary, the iterative reweighed *L*_2_ algorithm has the following schema:

#### Algorithm 1

Optimization algorithm

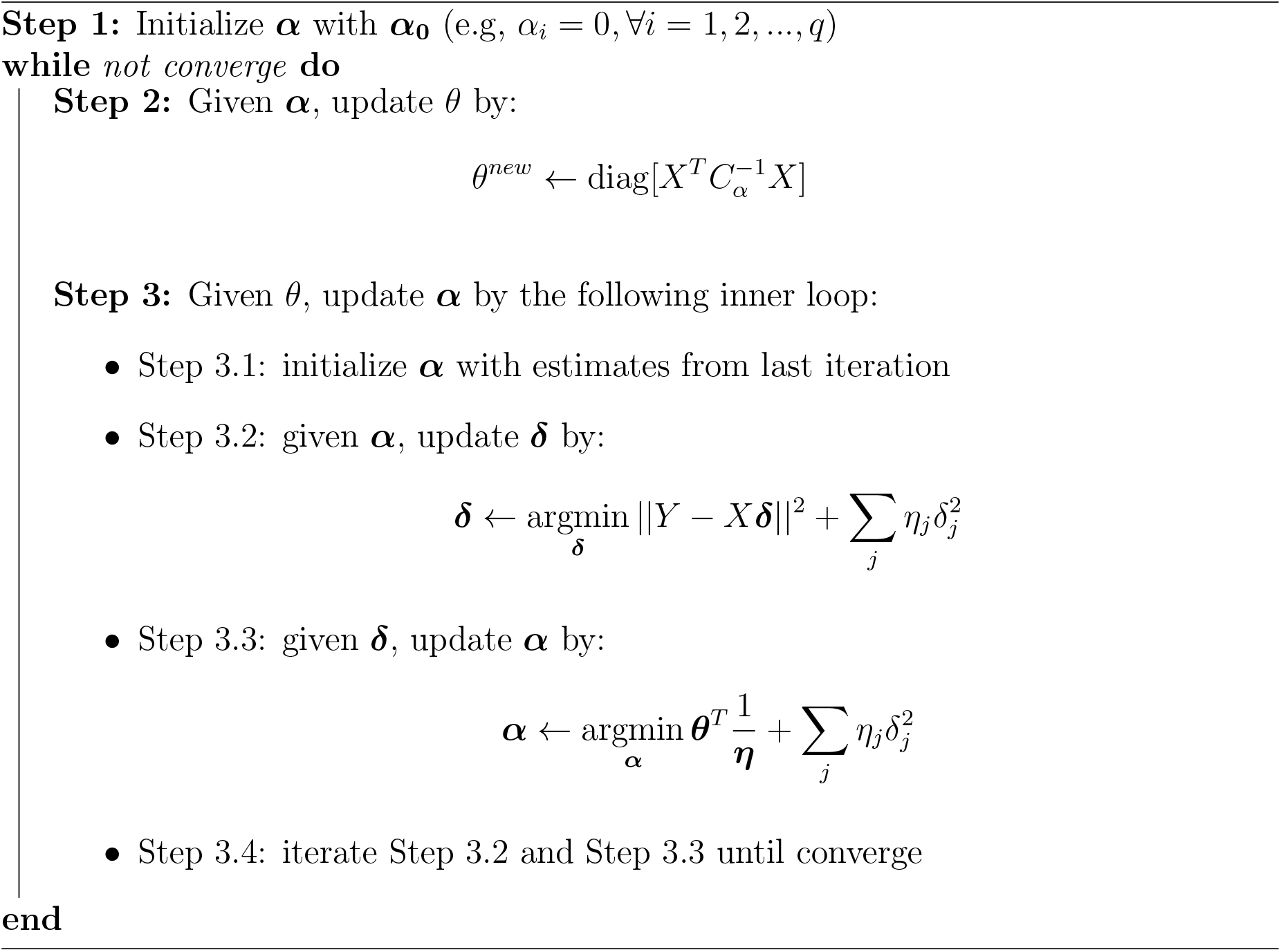

Once the hyperparameters ***α*** are estimated, and therefore the penalties known, the LASSO regression coefficients can be obtained using standard LASSO software (e.g. glmnet).

### 2.4 Extension to linear discriminant analysis for classification

So far we have been focused on LASSO linear model where the response variable is continuous. Here, following the scheme proposed by Mai *et al.* (2012) that builds high dimensional linear discriminate analysis (LDA) upon sparse penalized linear regression, we extend the xtune LASSO model to the framework of LDA with a binary response variable *Y* ∈ {1, 2}.

The LDA model assumes that *X* is normally distributed within each class, i.e,

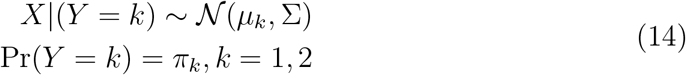

where *μ*_*k*_ is the mean of X within class *k* and Σ is the common within class covariance matrix. We adapt the following procedure proposed by Mai *et al.* (2012) to predict the class of *Y* based on xtune LASSO linear penalized regression:

Step 1. Let 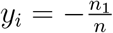 if *Y*_*i*_ = 1, and 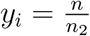 if *Y*_2_ = 2.
Step 2. Compute the solution to a penalized least squares problem:

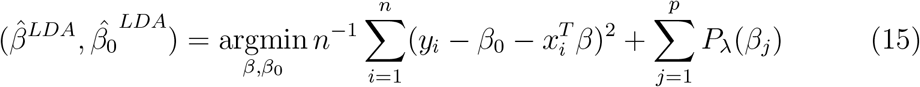
Step 3. Estimate the LDA model on the reduced data 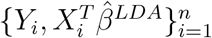. Assign observation *x* to class 2 if

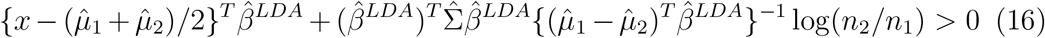

where 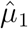, *n*_1_, 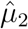, *n*_2_ are the sample mean vector and sample size within class 1 and class2, respectively. 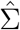 is the sample covariance matrix. *P*_*λ*_(.) is a generic sparsity-inducing regularization term, such as the single-penalty *L*_1_ norm, Elastic-Net or adaptive *L*_1_ norm in our case. We first solve (15) using xtune LASSO, then predict response variable class by (16).

## 3 Results

### 3.1 Simulation Studies

#### 3.1.1 Simulation setting

We performed a simulation study to evaluate the performance of the xtune LASSO under a range of scenarios obtained by varying the following key simulation parameters:

1. The true ability of the features to predict the outcome as measured by the signal to noise ratio (SNR), defined as Var(*X**β***)/*σ*^2^.
2. The informativeness of the external metadata, which is controlled by the number of non-zero hyperparameters *α*.
3. The number of predictor features *p*.
4. The proportion of predictive features, captured by the number of non-zero true regression coefficients ***β***.
5. The overall magnitude of the non-zero entries of the true regression coefficient vector ***β***, which is controlled by *α*_0_.
6. The overall degree of correlation between features.

The simulation data was generated according to the following steps: 1) set the parameter ***α*** controlling the informativeness of the external metadata; 2) generate the external metadata matrix *Z*; 3) generate the vector of regression coefficients ***β*** based on ***α*** and *Z*; 4) generate the feature data *X*; 5) generate the outcome *Y* based on *β*, *X* and an independent drawn random error.

The external metadata matrix *Z* of dimension *p* × *q* was generated to have 0-1 entries indicating a grouping of features. Specifically, entry *Z*_*jk*_ indicates whether the *j*^*th*^ feature belongs to group *k* (1) or not (0). The entries were independently generated to be zero or one with probability 0.2 and 0.8, respectively. A consequence of how *Z* was generated is that features can belong to more than one group, i.e. the groups can overlap. A column of 1 was appended to *Z* corresponding to the intercept *α*_0_, which controls the overall amount of shrinkage; the higher *α*_0_; the smaller the regression coefficients.

The true regression coefficient *β*_*j*_ were generated by sampling from a double exponential distribution with local parameter *μ* = 0 and scale parameter *b* = exp(*Z*_*j*_***α***). Threshold as described in Reid *et al.* (2013) was then applied to achieve sparsity. Specifically, the [*n^δ^*] largest *β*s in magnitude were retained, while all smaller entries were set to 0. The parameter *δ* controls the sparsity of the final ***β*** vector, with a lower value of *δ* inducing a sparser vector. The feature matrix *X* was simulated by sampling from a multivariate normal distribution with mean vector zero and covariance matrix Σ_*i,j*_ = *ρ*^|*i*−*j*|^.

Finally, for each simulation replicate, an independent error term *ϵ* was sampled from a normal distribution with mean 0 and a variance corresponding to the pre-set SNR. The outcome was generated as *Y* = *X****β*** + *ϵ*. Each simulated data is split into a training set (*n* = 200) and a test set (*n* = 1000). The performance of the xtune LASSO was assessed by the prediction *R*^2^ computed on the test using the model fitted in the training set. The large test set guarantees that the generalization/test error was accurately estimated. Penalty parameter tuning for the standard LASSO, which was fitted to compare with the proposed xtune LASSO, was performed by 10-fold cross-validation. One hundred replicates were generated for each scenario.

We consider nine scenarios varying each of the main simulation parameters described above:

1. *SNR* = 1, 3, 5
2. *p* = 500, 1000, 3000
3. External data informativeness: low, medium, and high. We fixed the sub-set of the ***α*** = (*α*_0_, −1, −0.78, −0.56, −0.33, −0.11, 0.11, 0.33, 0.56, 0.78, 1). The low external information is simulation from 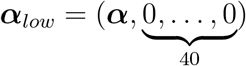. The medium external information is simulated from 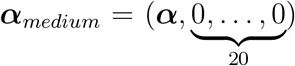, and the high external information is simulated from ***α***_*high*_ = ***α***. The idea is that non-informative external information has many noise variables.
4. *α*_0_ = 1, 3, 5
5. *δ* = 0.3, 0.5, 0.7
6. Different correlation magnitude *ρ* = 0, 0.3, 0.6, 0.9

The simulation results are summarized in Figure 1.

**Figure 1:**
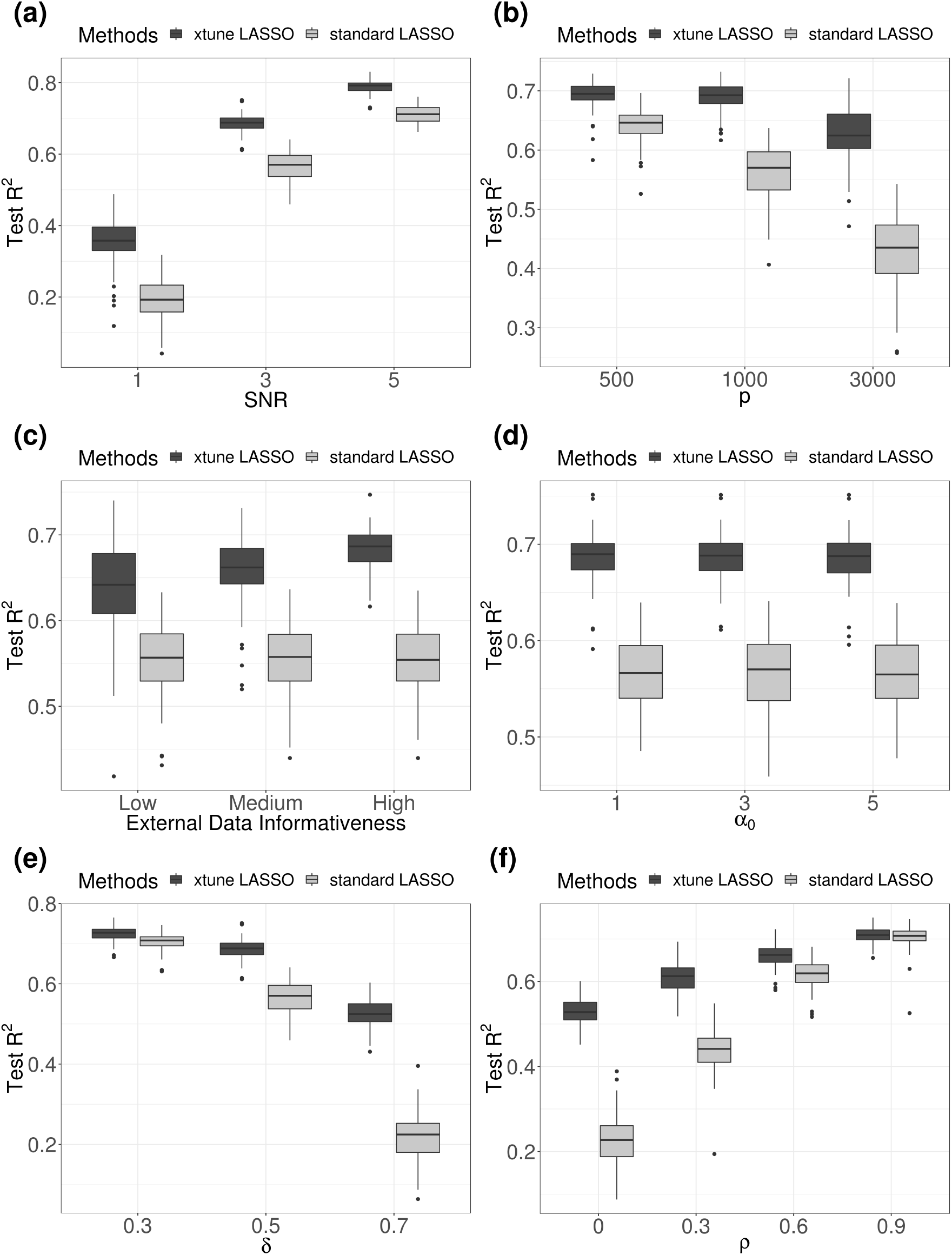
Simulation Results. Subplot (a): SNR = 1, 3, 5, with *n* = 200, *p* = 1000, *q* = 10, *α*_0_ = 3, *δ* = 0.5, *ρ* = 0.2. Subplot (b): *p* = 500, 1000, 3000, with *n* = 200, SNR = 3, *q* = 10, *α*_0_ = 3, *δ* = 0.5, *ρ* = 0.2. Subplot (c): *q* = 10, 30, 50, with *n* = 200, *p* = 1000, SNR = 3, *α*_0_ = 3, *δ* = 0.5, *ρ* = 0.2. Subplot (d): regression coefficients sparsity *δ* = 0.3, 0.5, 0.7, with *n* = 200, *p* = 1000, SNR = 3, *q* = 10, *α*_0_ = 3, *ρ* = 0.2. Subplot (e): overall penalty magnitude *α*_0_ = 1, 3, 5, *n* = 200, *p* = 1000, SNR = 3, *q* = 10, *δ* = 0.5, *ρ* = 0.2. Subplot (f): *ρ* = 0, 0.3, 0.6, 0.9, with *n* = 200, *p* = 1000, SNR = 3, *q* = 10, *α*_0_ = 3, *δ* = 0.5.

#### 3.1.2 Simulation results and data application of EB tuning

Figure 1 (a) shows box plots (across simulation replicates) of the prediction *R*^2^ for the standard and xtune LASSO as a function of the SNR of the metadata. As the SNR the prediction performance increases for both methods but the xtune LASSO has a better prediction across all levels of SNR considered. When the *p*/*n* ratio gets higher, both methods have decreased performance. However, xtune LASSO has a slower decreasing rate. When the *p*/*n* ratio is very high, we see that the performance of standard LASSO decreased dramatically and xtune LASSO significantly better prediction performance than standard LASSO. The *R*^2^ for standard LASSO remain the same across different level of external data informativeness. The performance of xtune LASSO decreases when the external information has low informativeness. In the case when the external information is completely useless, xtune LASSO performs slightly worse than standard LASSO (not shown).

Figure 1 (d) shows the effect of decreased ***β*** sparsity. The higher value of *δ*; the less sparse the model and the more non-zero regression coefficients. The value of xtune LASSO becomes more apparent when the model is less sparse. With *δ* = 0.5, 14 out of 1000 features are non-zero. Both methods have a decreased performance when the model has more signal features, but xtune LASSO has a slower decreasing rate. *α*_0_ controls the overall amount of shrinkage. From Figure 1 (e), the overall amount of shrinkage seems to have little effect on the performance of both methods. Notice that both methods have a higher prediction ability when the signal features (non-zero) features are correlated with each other. This is especially true for standard LASSO. The improved prediction of xtune LASSO over standard LASSO becomes smaller as the correlation between signal features *ρ*_*signal*_ increases. LASSO is known to have a decreased ability in variable selection when the features are highly correlated. It tends to select one variable from each highly correlated group and ignoring the remaining ones. (Hebiri and Lederer, 2013) studies the influence of correlation on LASSO prediction, and they suggest that correlation in features is not problematic for LASSO prediction. They also find that the prediction errors are mostly smaller for the correlated settings in experimental studies.

In summary, our simulation results show that the LASSO with individual penalties informed by meta-features can outperform the standard LASSO in terms of prediction when 1) the meta-features are informative for the regression effect sizes, 2) the true model is less sparse, and 3) the SNR is relatively high.

### 3.2 Applications

We exemplify the method’s performance on real data by considering two applications to clinical outcomes prediction using gene expression data.

#### 3.2.1 Bone density data

In the first example, bone biopsies from 84 women were profiled using an expression microarray to study the relationship between bone mass density (BMD) and gene expression. The goal is to predict the total bone density based on the gene expression profiles. The bone density is measured by the hip T-score derived from the women’s biopsies, with a higher score indicating higher bone density. The microarray data contains over 22,815 probe sets. The data were normalized using the RMA method as described in (Tharmaratnam *et al.*, 2016). Gene expression levels were analyzed on a logarithmic-2 scale. The bone density dataset is publicly available from the European Bioinformatics Institute (EMBL-EBI) Array Expression repository ID E-MEXP-1618.

Insights from previous study results are used as external covariates. (Reppe *et al.*, 2010) identified eight genes that are highly associated with bone density variation. We create a two-column external information *Z*. The first column is a column of all 1, representing the overall amount of shrinkage. The second column is a binary variable indicating whether each gene is one of the eight genes identified.

To illustrate the advantage of incorporating external information, we compare our proposed method to the standard LASSO and also the adaptive LASSO. The adaptive LASSO also adjusts the penalty individually for each coefficient, but the adaptive weight is obtained from the initial estimate of the coefficients using the same data rather than external information. Our implementation of adaptive LASSO utilizes the *adalasso()* function in the *parcor* R package.

The data were randomly split into a training data consisting of 80% of the observations and a test data set consisting of 20% of the observations. We fitted the adaptive LASSO, standard LASSO and our proposed method in the training data and evaluated their prediction performance in the testing data. We repeated 100 random splits of the full data into training and test sets. Figure 2 shows the MSE, *R*^2^ and the number of selected (non-zero) expression features across the 100 splits.

**Figure 2:**
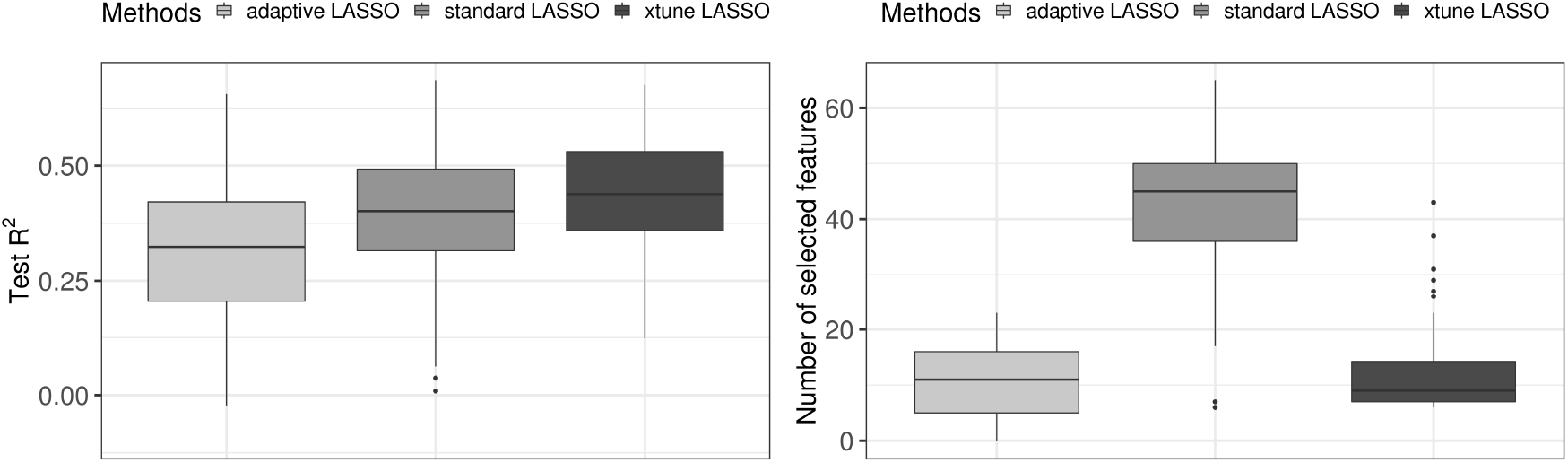
Compare prediction MSE, *R*^2^ and number of selected covariates of adaptive LASSO, standard LASSO and xtune LASSO using bone density data. The median *R*^2^ is 0.27 for adaptive LASSO; 0.38 for the standard LASSO, and 0.43 for xtune LASSO. The median number of selected covariates is 10 for adaptive LASSO, 43 for standard LASSO and 12 for the xtune LASSO.

Overall, we see that the externally tuned LASSO has better prediction performance than the standard LASSO while selecting a more parsimonious model. The adaptive LASSO does not perform well in this data example. To gain further insight into the prediction performance results, we examined the penalties applied to the regression coefficients by each of the methods when fitted on the full data. The tuning parameter chosen by standard LASSO using cross-validation is 0.16, while for the xtune LASSO the estimated tuning parameter is 0.26 for the gene expression features without external information and 0.016 for the expression features with external information, resulting in larger effects estimates for the latter group.

#### 3.2.2 Breast cancer data

In the second example, we apply the xtune LASSO to a breast cancer dataset. The data is from the Molecular Taxonomy of Breast Cancer International Consortium (METABRIC) cohort (Curtis *et al.*, 2012) (https://ega-archive.org/dacs/EGAC00001000484). 29,476 gene expression profiles and three clinical variables (age at diagnosis, progesterone receptor status and lymph node status) were used for the prediction of five-year survival (survived or died). Patients followed-up for less than five years with no record of mortality were excluded from the analysis. We used a subset of the METABRIC data with patients that are Estrogen receptor (ER) positive and human epidermal growth factor receptor 2 (HER2) status negative. The data contains a discovery set of 594 observations and an additional validation set of 564 observations. The models were trained in the discovery set and tested on the validation set.

The external information used for the xtune LASSO model is based on the results of (Cheng *et al.*, 2013), where groups of genes referred to as ‘metagenes’ that are prognostic in all cancers including breast cancer were identified. In specific, (Cheng *et al.*, 2013) analyzed six gene expression datasets from different cancer types and present three multi-cancer attractors with strong phenotypic associations: a lymphocyte-specific attractor (LYM), a mesenchymal transition attractor strongly associated with tumor stage (MES), and a mitotic chromosomal instability attractor strongly associated with tumor grade (CIN). The LYM, MES, and CIN metagenes consist of 169, 134 and 108 genes, respectively. Therefore, the meta-feature matrix ***Z*** consists of four external covariates: the first covariates is a binary variable indicating whether a predictor in *X* is a clinical feature (1) or a gene expression feature (0) and the next three columns map genes that belong to with LYM, MES, and CIN respectively.

We compared the xtune LASSO incorporating the meta-features described above, with the standard and the adaptive LASSO. As in the first example, standard LASSO was tuned by repeated 10 fold cross-validation and the adaptive LASSO is implemented using the *adalasso()* function in the *parcor* R package.

Table 1 compares the AUC, the number of selected features and the computation time for the standard, the adaptive and the xtune LASSO. Figure 3 shows the receiver operating characteristic (ROC) curves for the three methods. The xtune LASSO has the best prediction performance among all three methods. The three clinical variables are given a very small penalty in xtune LASSO and are both estimated to be non-zero. The penalty chosen by repeated cross-validation for the standard LASSO is 0.020. For xtune LASSO, the penalty applied to clinical variables, ‘non-attractor’ genes, LYM metagenes, MES metagenes and CIN metagenes is 0.001, 0.114, 0.030, 0.038 and 0.028, respectively. This illustrates how the xtune LASSO can induce differential shrinkage among the features according to their empirical importance. In this example, xtune LASSO shrinks the coefficients corresponding to expression features that do not belong to any metagene (the vast majority) toward zero much more aggressively than the standard LASSO, while it shrinks the clinical variables and the expression features in metagenes LYM and MES much less than the standard LASSO. The standard LASSO shrink the coefficients for the two clinical variables to 0, effectively selecting them out of the model. Adaptive LASSO selected out one of the clinical variables (lymph node status). In agreement with our simulation results, the xtune LASSO also yielded a much more parsimonious model with only 10 selected features while the standard LASSO selected 207 features. However, the xtune LASSO is more computationally demanding than the standard LASSO tuned by cross-validation.

**Table 1:** Compare AUC, number of selected covariates and computation time (in minutes) for standard LASSO, adaptive LASSO and xtune LASSO

|  | Standard LASSO | Adaptive LASSO | xtune LASSO |
| --- | --- | --- | --- |
| AUC | 0.675 | 0.718 | 0.746 |
| No. of selected covariates | 207 | 5 | 10 |
| Computation time | 1.58 | 30.81 | 12.15 |

**Figure 3:**
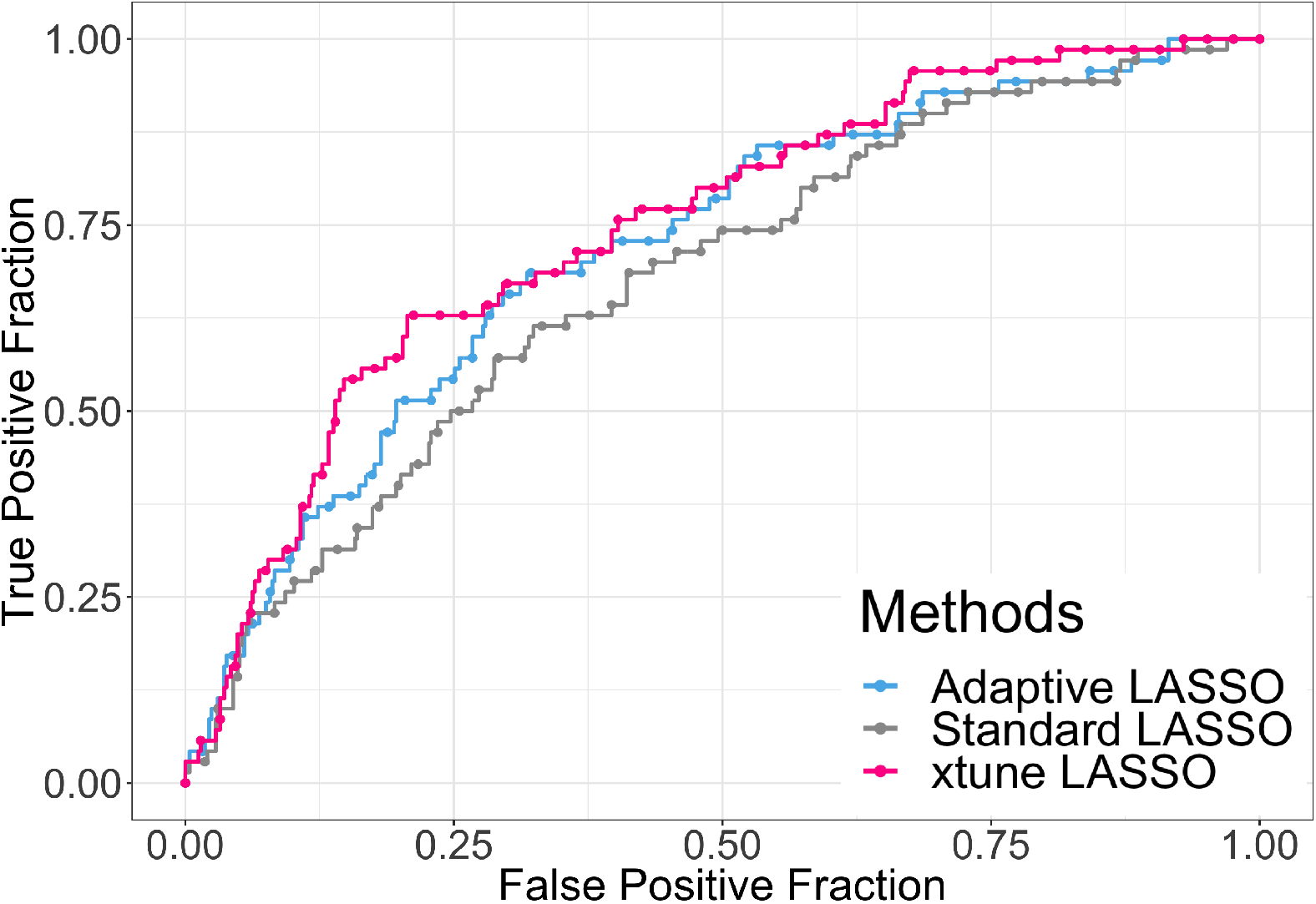
ROC curves for adaptive LASSO, standard LASSO and xtune LASSO applied on the breast cancer dataset

## 4 Discussion

We introduced xtune LASSO, a new approach implemented in the namesake R package (Zeng and Lewinger, 2019) for integrating external meta-features to inform the tuning of penalty parameters in LASSO regression. We showed through simulation and real data examples that xtune can yield better prediction performance than standard LASSO. These findings are consistent with the work of (van de Wiel *et al.*, 2016) who proposed a method for estimating group-specific penalties for ridge regression and showed that the use of prior knowledge can improve prediction performance.

xtune LASSO differs from related methods (Pan *et al.*, 2010), (Boulesteix *et al.*, 2017), (Liu *et al.*, 2018) in our use of an empirical Bayes approach to estimate the penalty parameters, rather than relying on cross-validation. In the particular case of no external meta-features (i.e. *Z* is a vector of 1s) xtune performs empirical Bayes tuning of the single LASSO penalty parameter, providing an alternative to standard tuning by cross-validation. However, cross-validation becomes impractical with more than a handful of penalty parameters, while empirical Bayes tuning allows xtune to handle a much larger number of individualized penalties.

In both our real data application examples, the meta-features are categorical variables that group features into subsets. A categorical meta-feature also arises when the features originate from different data types (e.g. gene expression, methylation, somatic mutations). (Liu *et al.*, 2018) showed that having separate penalty parameters for each data type can yield better prediction performance. However, xtune is not limited to categorical meta-features and can equally handle quantitative ones, such as effect estimates or p-values for the association of the features and the outcome of interest or related outcomes, which can be exploited to integrate information from previous studies.

Although prediction performance has been our main focus of interest, our results also show that for the range of simulation scenarios we considered and in the two real data applications, xtune tends to yields sparser and therefore more interpretable models than standard LASSO regression. Moreover, the estimates of the metafeature coefficients ***α*** can yield additional biological insights as they capture the importance of the meta-features for predicting the outcome.

However, we did note that when the number of meta-features *q* relative to the sample size *n* is large, the ***α*** estimates may not be stable. A related limitation is that in its current implementation xtune does not scale to ultra high dimensional datasets. Typical datasets that xtune LASSO can currently handle have sample size of up to *n* 5000, with *p* 50, 000 features and *q* 100 meta-features. However, we believe that future algorithmic improvements along with parallel computing can extend the applicability of xtune to larger datasets and larger numbers of meta-features. To further widen the range of applicability of xtune, we are pursuing extensions to binary (logistic regression) and time to event (Cox regression) outcomes, as well as the incorporation of the Ridge and Elastic-Net penalties in addition to the LASSO.

